# Analysis of alternative polyadenylation from long-read or short-read RNA-seq with LAPA

**DOI:** 10.1101/2022.11.08.515683

**Authors:** Muhammed Hasan Çelik, Ali Mortazavi

## Abstract

**Motivation:** Alternative polyadenylation (APA) is a major mechanism that increases transcriptional diversity and regulates mRNA abundance. Existing computational tools to analyze APA have low precision because these tools are designed for short-read RNA-seq, which is a suboptimal data source to study APA. Long-read RNA-seq (LR-RNA-seq) accurately detects complete transcript isoforms with poly(A)-tails, providing an ideal data source to study APA. However, current computational tools are incompatible with LR-RNA-seq.

**Results:** Here, we introduce LAPA, a computational toolkit to study alternative polyadenylation (APA) from diverse data sources such as LR-RNA-seq and short-read 3’ sequencing (3’-seq). LAPA counts and clusters reads with poly(A)-tail, then performs peak-calling to detect poly(A)-site in a data source agnostic manner. The resulting peaks are annotated based on genomics features and regulatory sequence elements such as presence of a poly(A)-signal. Finally, LAPA can perform robust statistical testing and multiple testing correction to detect differential APA.

We analyzed ENCODE LR-RNA-seq data from human WTC11, mouse C2C12 myoblast, and C2C12-derived differentiated myotube cells using LAPA. Comparing LR-RNA-seq from different platforms and library preparation methods against 3’-seq shows that LR-RNA-seq detects poly(A)-sites with a performance of 75% precision at 57% recall. Moreover, LAPA consistently improved TES validation by at least 25% over the baseline transcriptome annotation generated by TALON, independent of protocol or platform. Differential APA analysis detected 788 statistically significant genes with unique polyadenylation signatures between undifferentiated myoblast and differentiated myotube cells. Among these genes, 3’ UTR elongation is significantly associated with higher expression, while shortening is linked with lower expression. This analysis reveals a link between cell state/identity and APA. Overall, our results show that LR-RNA-seq is a reliable data source for the study of post-transcriptional regulation by providing precise information about alternative polyadenylation.

**Availability:** LAPA is publicly available at https://github.com/mortazavilab/lapa and PyPI.

## 1 Introduction

Alternative polyadenylation (APA) is a post-regulatory process resulting in heterogeneous transcripts with unique 3’ end sites. During polyadenylation, RNAs are cleaved at a specific site called poly(A)-site where a 50-100 nucleotide (nt) poly(A)-tail is appended to the 3’ end of each transcript [6]. Poly(A)-sites are usually located in the 3’ UTR and defined by regulatory sequence elements in the vicinity, such as the poly(A) motif (“AATAAA”). 70% of mRNA encoding genes have more than one poly(A)-site [31]. Minor non-canonical poly(A)-sites also exist in exonic and intronic regions. Poly(A)-sites that result in a longer 3’ UTR are called a distal poly(A)-site, while poly(A)-sites resulting in a shorter 3’ UTR are called a proximal poly(A)-site.

The 3’ UTR plays a critical role in post-transcriptional regulation because the 3’ UTR contains targets for microRNA (miRNAs) and RNA binding proteins (RBPs). Elongation of 3’ UTR length during evolution from a median length of 140 base pair (bp) in worms to 1-2 kilo bp in humans suggests an increase in both the complexity of post-transcriptional regulation and the importance of regulatory elements in the 3’ UTR region [23]. 3’ UTR isoforms resulting from specific proximal or distal poly(A)-site usage can lead to differential exclusion or inclusion of regulatory sequence elements such as RBP binding sites [31]. Distal poly(A)-site usage can lead to 3’ UTR elongation, resulting in miRNA-binding site gain and reduced gene expression. Conversely, proximal poly(A)-site usage can lead to 3’ UTR shortening, resulting in miRNA-binding site loss. As a result, APA regulates post-transcriptional processes such as mRNA folding, stability, localization, and translational efficiency [22]. Thus, APA is essential for cell state and identity. For example, cells have a unique APA signature during proliferation and differentiation [12, 31]. Consequently, the misregulation of APA is associated with genetic diseases and cancer [12].

Definitive evidence of polyadenylation can be obtained by sequencing the poly(A)-tail of mRNA. There are many computational tools that detect APA using short-read RNA sequencing [7]. A systematic benchmark of these tools demonstrates that tools based on short-read RNA-seq suffer from poor precision and recall [7]. Another systematic benchmark study [28] shows that the true positive rate (TPR) of short-read RNA-seq based APA tools is below 60% at any false positive rate (FPR) when compared against long-read RNA-seq ground truth. The experimental limitations of short-read RNA-seq can explain the poor performance of these tools. Only a small subset of reads are poly(A)-tailed in short-read RNA-seq due to coverage bias at 5’ and 3’ of the transcripts (Figure-1.a). Thus, solely relying on these poly(A)-tailed reads is not feasible. Alternative strategies, such as detecting sharp changes in read coverage or leveraging prior annotation of poly(A)-sites, are also employed by short-read RNA-seq based APA tools. Nevertheless, these approaches have their own issues. For example, technical biases can create noise in the form of fluctuations in short RNA-seq coverage, and relying on prior annotation poly(A)-sites cannot detect novel poly(A)-sites [7].

3’ sequencing (3’-seq) is a common name for a set of purpose-specific short-read based protocols that detects APA. 3’-seq is enriched for reads containing poly(A)-tail to overcome the limitation of standard RNA-seq (Figure-1.a). More than fifteen 3’-seq [8] sequencing protocols were proposed, such as QuantSeq3 and 3’ READS. Despite advances on the experimental side, there are no user-friendly software packages that can analyze 3’-seq data in a protocol-independent manner. Usually, 3’-seq data is analyzed with custom scripts that are not generic enough to apply to other protocols. The lack of a publicly-available computational toolkit for 3’-seq creates a bottleneck to studying APA. Additionally, 3’-seq data are not as abundant as standard short-read RNA-seq given that these protocols are purpose-specific.

An alternative data source to study APA is long-read RNA sequencing (LR-RNA-seq). LR-RNA-seq facilitates the study of APA because LR-RNA-seq reads can capture complete transcript isoforms with poly(A)-tails (Figure-1.a) [30, 4, 3, 28]. Although existing LR-RNA-seq tools can detect reads with poly(A)-tails, those tools are either mainly focused poly(A)-tail length estimation [16, 20] or do not provide a software package to analyze APA [28]. To our best knowledge, there is no user-friendly software for clustering and peak calling poly(A)-sites or analysis of APA from LR-RNA-seq. Also, major LR-RNA-seq protocols/platforms have not been benchmarked in-depth for the purpose of APA. To advance the study of alternative polyadenylation, we developed LAPA, a computational toolkit to study APA from diverse data sources such as LR-RNA-seq and 3’-seq. LAPA is available in PyPI and provides a user-friendly API to analyze APA given an alignment file. LAPA is generic enough to analyze any 3’-seq or long-read RNA-seq protocol. In this paper, we benchmark the performance of a range of long-read platforms and libraries to detect alternative polyadenylation using LAPA. Moreover, we demonstrate downstream applications of LAPA, such as correction of annotated transcript ends and differential APA analysis in myogenesis.

## 2 Methods

LAPA is a highly modular (Figure 1-b) software that takes alignment (BAM) files as input and annotates APA after platform/protocols-specific preprocessing (Supplementary Method-1). The output file structure of LAPA and the content of files are described in Supplementary Method-3. We provide both Python and user-friendly command-line (CLI) interfaces to our users. The modules described below can be repurposed for applications beyond APA.

**Figure 1:**
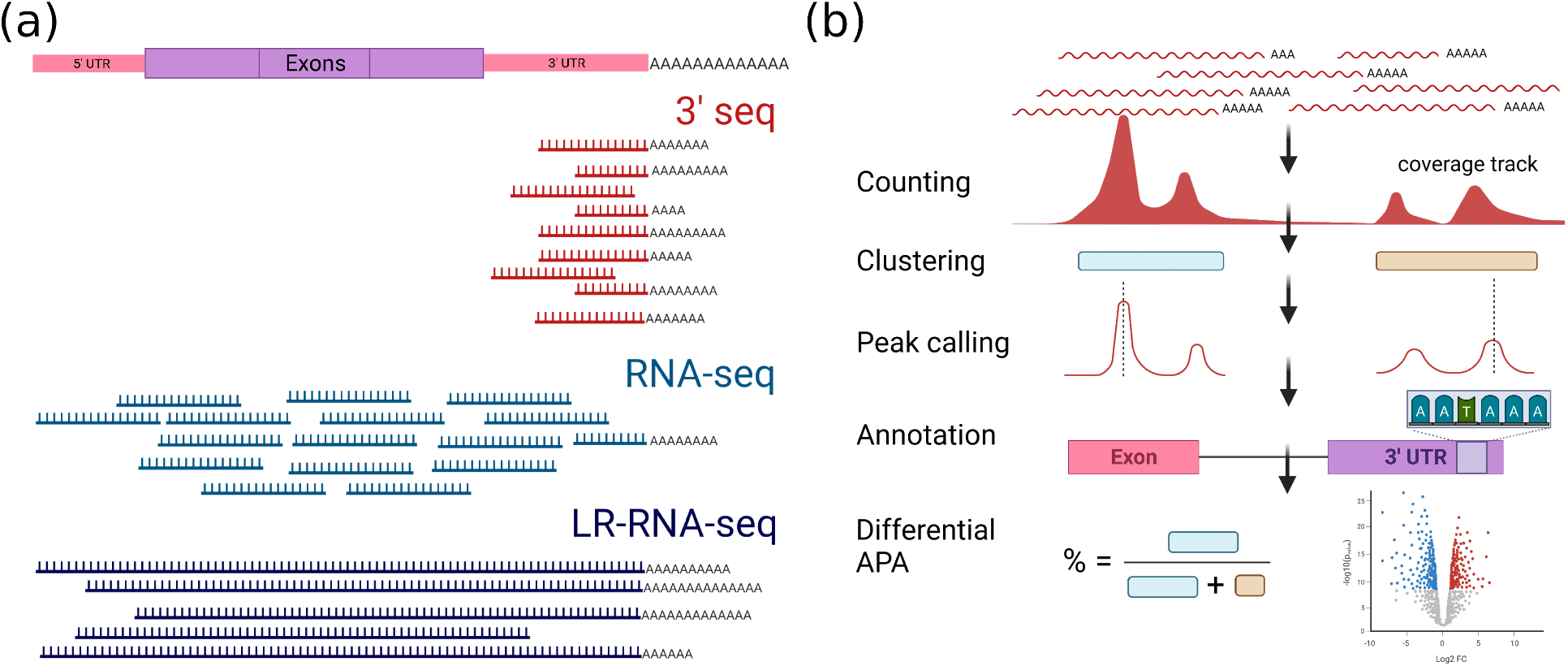
(a) Major sequencing assay to study alternative polyadenylation: 3’-seq (blue), short-reads (red) and long-reads (purple). (b) Computational steps of LAPA.

### 2.1 Read counting

The first step of LAPA is counting 3’ ends of the reads to create read-end counts per position with respect to the reference genome. Low-quality reads (default mapq>10) and secondary alignments are filtered during the counting process. The algorithmic complexity of counting is linear in terms of the number of reads. Additionally, we calculate the total coverage of positions where read-end coverage is non-zero. Total coverage is defined by the number of reads ending or covering a specific position. Based on those two coverage tracks, we further calculate the read-end signal, indicating the percentage of reads ending at a specific position which is calculated by dividing read-end counts to total coverage. We store all coverage tracks in sparse bigwig file format (Supplementary Method-3) for visualization in the genome browser and further processing.

During the counting step, LAPA detects reads with poly(A)-tail if poly(A)-tails are present in the alignment file. Poly(A)-tailed reads contain the sequence of ‘A’ base pairs at the 3’ end, and this homopolymeric ‘A’ sequence does not align to the reference genome sequence but rather is soft-trimmed in the alignment file. Aligners usually align poly(A)-tails to reference the genome and shift potential poly(A)-site if there are corresponding sequences of ‘A’ bp pairs such as internal priming sites in the reference genome sequence. We detect and correct such misaligned poly(A)-tail bases. Based on the detected poly(A)-tails, LAPA provides the distribution of poly(A)-tail lengths obtained in this step as a quality control measure. LAPA provides an optional filter that includes reads based on the presence of poly(A)-tail with a certain length. This option is not recommended for data from 3’-seq protocols, which typically only yield reads with short poly(A) tails (<20 bp) or reads with no poly(A) tails at all. Poly(A)-tail filtering is well-suited for LR-RNA-seq if most of the reads have relatively long poly(A)-tails (>20 bp). However, poly(A)-tails in the LR-RNA-seq reads are commonly trimmed off during preprocessing for both Oxford Nanopore (ONT) and PacBio (Supplementary Method-2). Thus, poly(A)-tail counting requires re-processing and alignment of raw data, which is not user-friendly and computationally expensive. Therefore, we implemented our counting strategy in a way that also works without poly(A)-tailed reads. Although there can be partial reads in LR-RNA-seq with incomplete 5’-3’ ends, those reads are unlikely to cluster together unless there is internal priming; hence, partial reads will be filtered out in downstream steps.

### 2.2 Poly(A)-site clustering

We create poly(A)-clusters from the read-end counts and coverage obtained from the previous steps. In clustering, we scan a chromosome from start to end, iterate over the read-end counts, and initialize a cluster if the read-end count in a specific position is higher than the cutoff of x% of the total coverage and more than N reads (default x=5% and N=3). The read number cutoff is further tuned based on the replication rate (Method-2.5). Poly(A)-clusters are extended as long as read-end counts exceed the cutoff and terminated based on patience parameters where read-end counts are consistently below the cutoff for N base pairs (default N=25 bp). The algorithmic complexity poly(A)-site clustering is linear in unique genomic positions with a non-zero end count.

### 2.3 Peak calling

The previous detection of a poly(A)-site requires calling the position with the peak number of read-end counts. However, the number of read-ends may fluctuate in a poly(A)-cluster; thus, read-end counts are smoothed with the moving average of Gaussian kernel with a window size of 5 and standard deviation of 1 provided as:

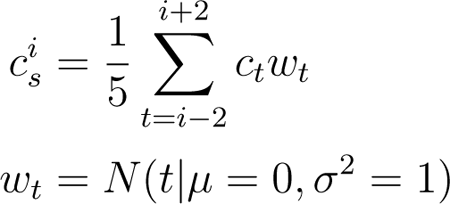

where *c*_*t*_ is the read count at position *t, C^i^_s_* is the smoothed read count and *w*_*t*_ is weight obtained from the Gaussian kernel. After the smoothing, the position with the maximum smoothed count (*C^i^_s_*) is selected as the polyadenylation site.

### 2.4 Cluster annotation

We annotate each poly(A)-site as exonic, intronic, 3’ UTR, etc., based on the genomic features from a standard genome annotation. The number of poly(A)-sites per genomic feature provides quality control for the data, given that most of the poly(A)-sites are expected to appear in the 3’ UTR region. Moreover, the poly(A)-signal (Figure-3.b) is also expected to appear in the vicinity of the polyadenylation site. Hence, we search for the canonical alternative polyadenylation motif (AATAAA) up to 10 bp upstream and 60 downstream of poly(A)-site [13].

**Figure 2:**
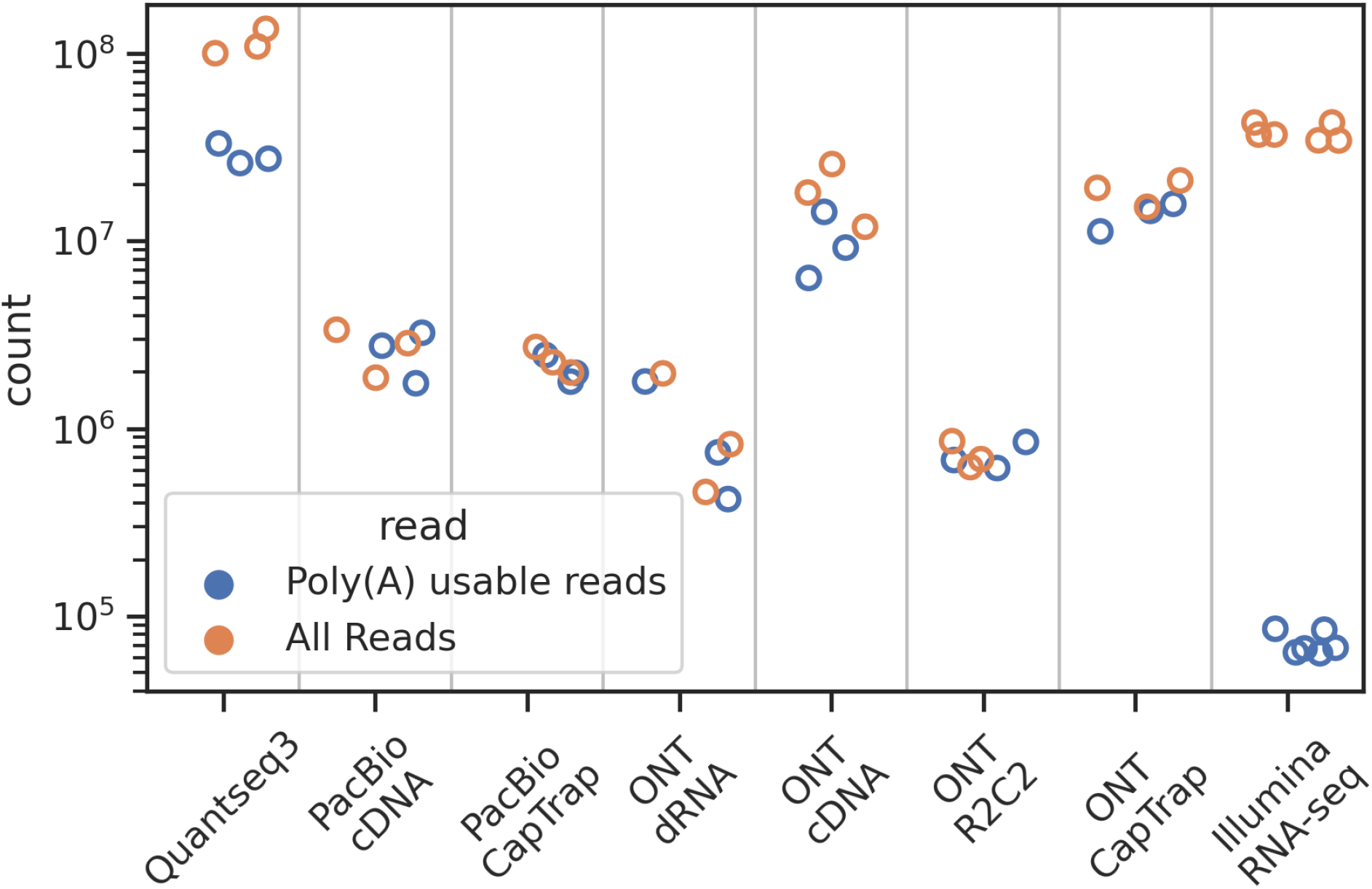
Number of total read yields from each sequencing platform/protocol and the number of reads usable to detect alternative polyadenylation.

**Figure 3:**
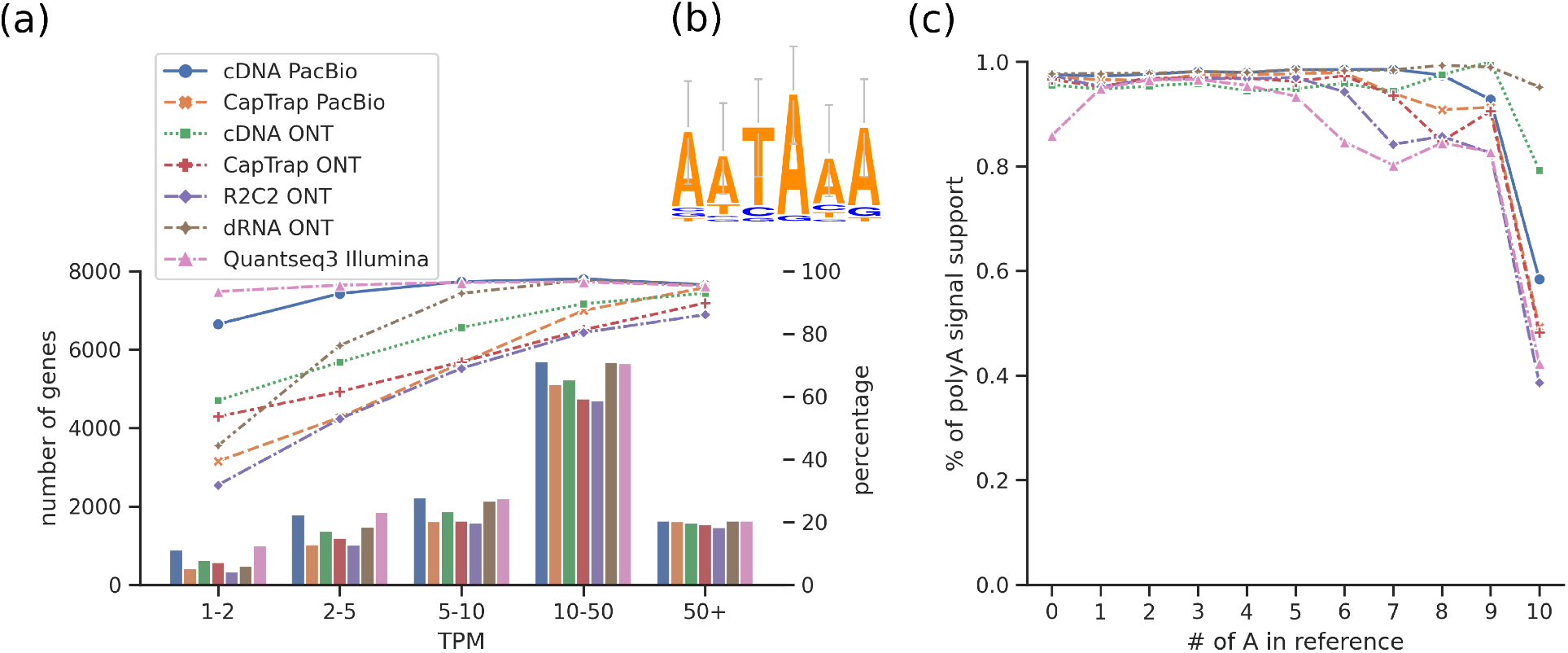
(a) Percentage of poly(A)-signal support of the major poly(A)-site of genes at different TPM levels for each sequencing platform/protocol. (b) Sequence logo of poly(A)-signal motif (c) Depletion of poly(A)-signal support of poly(A)-clusters with increasing A bp in the reference genome sequence following poly(A)-site due to internal priming.

Internal priming sites are a segment of DNA containing consecutive A base pairs mimicking a poly(A)-tail. Internal priming introduces false-positive poly(A)-sites. Thus, we count a number of A bp in the reference genome sequence following poly(A)-sites. If the 10 bp following a poly(A)-site contains more than 7 bp of A and the poly(A)-signal is missing in the vicinity, LAPA filters out the site because it is likely an internal priming site.

### 2.5 Replication rate

Poly(A)-clusters detected by one experiment may not be replicated by technical replicates due to issues like batch effects leading to false positive poly(A)-site detection. To limit the irreproducible discovery rate [19] of the poly(A)-cluster, we tune the threshold of the minimum number of reads based on the replication rate. Replication rate (*R*) is calculated with the following formula:

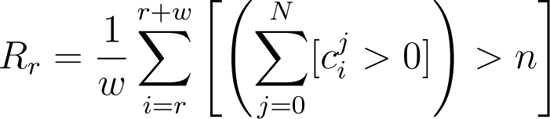

where clusters are first ranked (*r*) by read-end counts, cluster *c* is considered replicated if cluster observed in at least *n* of *N* samples (default *n* = 2); then the moving average of replication is calculated with a window size of *w* (default *w* = 1000). The cutoff for read-end counts is chosen to ensure a replication rate of at least *x*% (default x=95%). Moreover, the low replication rate of clusters with high read-end counts indicates a potential issue in the data source; thus, we provide replication rate statistics as quality control to users to spot such issues.

### 2.6 Statistical testing

To quantify the level of poly(A)-site usage, we define the *usage* metric indicating the percentage of use of a specific poly(A)-site *i* given all the poly(A)-sites in the gene. The usage metric is calculated by dividing the read-end counts of poly(A)-cluster by the total number of reads in the gene:

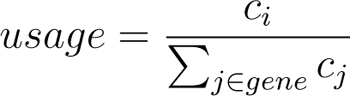

We implemented two statistical tests based on the count data: (1) the Fisher’s exact test, which compares two groups, and the (2) beta-binomial test, which compares multiple conditions with dispersion (Supplementary Method-4):

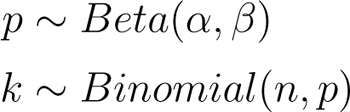

### 2.7 Improved transcript start and end annotation

Novel transcriptome annotations can be generated from LR-RNA-seq. The computational strategy we proposed in this paper is highly optimized for poly(A)-site detection. Thus, we investigated whether we can improve transcript end sites of transcriptome annotations generated using LR-RNA-seq. Since LAPA is modular, its modules can be repurposed for transcript start site (TSS) detection from long reads. We repurposed the counting module for read-start counting by counting 5’ ends rather than 3’ ends; then applied the clustering and peak calling steps out-of-the-box on read-start counts. As a result, we obtained TSSs from LR-RNA-seq. We spotted linking reads which start in a TSS-cluster and end in a poly(A)-cluster. Then, we obtained a splice chain of these linking reads using TALON [34]. Finally, we created a transcriptome annotation with improved TSS and TES sites and exported this improved annotation as GTF and abundance files, consistent with TALON file format.

### 3 Results

We analyzed LR-RNA-seq data from a range of protocols and platforms (cDNA and CapTrap for PacBio; dRNA, R2C2, and cDNA CapTrap for ONT) from LRGASP [25] with LAPA. We additionally used the short-read Quantseq3 (3’-seq) and single-ended Illumina WTC11 datasets part of LRGASP (Supplementary Method-1). Each protocol has at least three replicates. We run LAPA with its default arguments to avoid overfitting parameters for analyses.

### 3.1 Number of reads usable for poly(A)-site detection

We calculated the number of reads that can be utilized to detect APA (Figure-2, Supplementary Table-1). Firstly, there are approximately 105 ± 17 (sd) million raw Quantseq3 reads per replicate. Filtering reads for alignment quality and 18 bp of A required by the Quantseq3 protocol resulted in approximately 27 ± 3 million reads. There are 35 ± 4 million raw reads in the Illumina, yet, filtering reads with 18 bp poly(A)-tails yield 66,276 ± 9,391 reads per replicate. There are usually fewer reads at the start and end of transcripts, and most of the reads are intermediate because of the coverage bias of short-read RNA-seq. Therefore, we did not apply LAPA on short-read RNA-seq in downstream analysis because our approach is not well suited for short-read RNA-seq due to low poly(A)-tailed read numbers. LR-RNA-seq sequences complete transcript isoforms (Supplementary Figure-1, Supplementary Table-2). Thus, we used all aligned reads from LR-RNA-seq after quality filtering. LR-RNA-seq methods provide millions of reads that are usable to detect APA. Specifically, the PacBio platform provides 2.4 ± 0.7 million and 1.9 ± 0.3 million mapped reads with average read lengths of 2,525 and 1,042 for cDNA and CapTrap protocols, respectively. From the ONT platform, we obtained 9.1 ± 3.7 million cDNA and 12.7 ± 2.1 million CapTrap reads with average read lengths of 738 and 912 bp respectively.

Although the number of reads is higher in the ONT cDNA and CapTrap samples compared to the PacBio samples, the read length is shorter, suggesting the existence of partial reads. ONT dRNA and R2C2 with 0.9 ± 0.6 million and 0.6 ± 0.1 million reads respectively have the least number of reads across all the LR-RNA-seq samples. However, the dRNA protocol is strongly enriched for poly(A)-tailed reads because RNA adapters during chemistry are ligated onto the 3énds of poly(A)-tail, thus reads are always generated from the 3énd [11, 14]. Therefore, dRNA may provide an advantage for poly(A)-site detection by capturing reads with complete poly(A)-tails. Overall, LR-RNA-seq provides a sufficient number of reads to study APA, overcoming the limitation of short-RNA-seq.

We calculated reads ending in the proximity of transcript end sites (TES) from GENCODE [10]. The analysis indicates that reads result in very sharp and narrow (∼5 bp) peaks around the annotated sites (Supplementary Figure-2.a). This result shows that counting read ends can detect poly(A)-sites precisely. We provide this analysis as a quality control measure for LAPA users because the lack of sharp peaks around annotated sites indicated potential issues with the input data source. We further investigate the percentage of reads ending in the vicinity of TES annotated in GENCODE (Supplementary Figure-2.b). Only 34 ± 6% of the reads end in the vicinity of annotated TES regardless of protocol and platform (Supplementary Table-5). This indicates that there are many poly(A)-sites that are not annotated in GENCODE.

### 3.2 Poly(A)-Signal Support and Internal Priming

We calculated a number of genes with at least one poly(A)-cluster detected in the WTC11 cell line and plotted those genes based on gene expression (Figure-3.a). Then we investigated the number of genes where the major poly(A)-site has the poly(A)-signal of “AATAAA” (Figure-3.b). Canonical poly(A)-sites contain a poly(A)-signal, so looking for the poly(A)-signal in the vicinity of a major poly(A)-site provides a sanity check for poly(A)-cluster calls. 80% of major poly(A)-sites from highly expressed genes (10+ TPM) have poly(A)-signal support across all samples. Moreover, cDNA PacBio and ONT dRNA have over 90% support when gene expression is 5+ TPM. ONT R2C2 and PacBio/ONT CapTrap protocols have lower performance. R2C2 ONT has lower read numbers compared to other protocols. The CapTrap protocol is designed to compare the 5’ end of the transcripts, so it is not the preferred protocol to study the 3’ end of transcripts. As another check, we investigated and observed that 95 ± 4 % of major poly(A)-sites are located in the 3’ UTR of protein-coding genes across all samples and expression levels (Supplementary Table-4). Altogether, this analysis shows that LAPA can detect poly(A)-site with poly(A)-signal for expressed genes.

Internal priming is the primary driver of false-positive poly(A)-site discovery. Internal priming sites do not contain the sequence context of true poly(A)-sites. Thus, we investigated poly(A)-signal depletion with the increasing number of A base pairs in the reference genome sequence after the poly(A)-site (Figure-3.c). LAPA counts and reports the number of A base pairs in the reference genome sequence following 10 base pairs. Analysis based on these counts indicates a depletion of poly(A)-signal if more than 7 base pairs of A are observed in the reference genome sequence. The poly(A)-signal depletion is an indication of internal priming, and all the protocols except dRNA suffer from internal priming. Thus, we define poly(A)-sites without poly(A)-signal as internal priming sites. If poly(A)-sites have more than 7 base pairs A in the genome, those poly(A)-sites are filtered in LAPA.

### 3.3 Comparison of protocols and platforms

We compared the overlap of poly(A)-sites between different protocols and libraries. Firstly, we subset for poly(A)-sites from genes with >=5 TPM per sample, which represents a high confidence set. Then, we analyzed if other samples can detect this high confidence set of poly(A)-sites of a sample by measuring overlap. There is a high agreement of poly(A)-sites detected by LR-RNA-seq samples (Figure-4.b). For example, cDNA PacBio can detect 80-90% of the poly(A)-sites detected by other methods. Quantseq3 has the highest overlap by detecting most poly(A)-sites detected by other samples, yet other methods cannot detect all the poly(A)-sites detected by Quantseq3. This can be explained by the much higher read coverage of Quantseq3 compared to other methods (Supplementary Figure-4, Supplementary Table-1,6). Thus, we chose a high confidence set of Quantseq3 poly(A)-sites as ground truth and calculated the precision-recall curve as shown in figure-4.a (Supplementary Table-7). ONT dRNA and PacBio cDNA are the best performing methods, with the area under the precision-recall curve of 84% and 81%, respectively. Both of these methods have precision over 90% at the recall of 50%. Other methods have 60-80% precision at 50% recall. We further investigate the impact of different read-end counting settings on performance. Specifically, we asked if subsetting for the reads with at least 20 bp poly(A)-tail affects the performance. We observe that subsetting reads with poly(A)-tails have no impact on the performance (Supplementary Figure-5). The result shows that LAPA can detect poly(A)-sites even if poly(A)-tails are trimmed (Supplementary Figure-3, Supplementary Table-3). Overall, the benchmark shows that LAPA can detect poly(A)-sites with high precision and recall from LR-RNA-seq.

**Figure 4:**
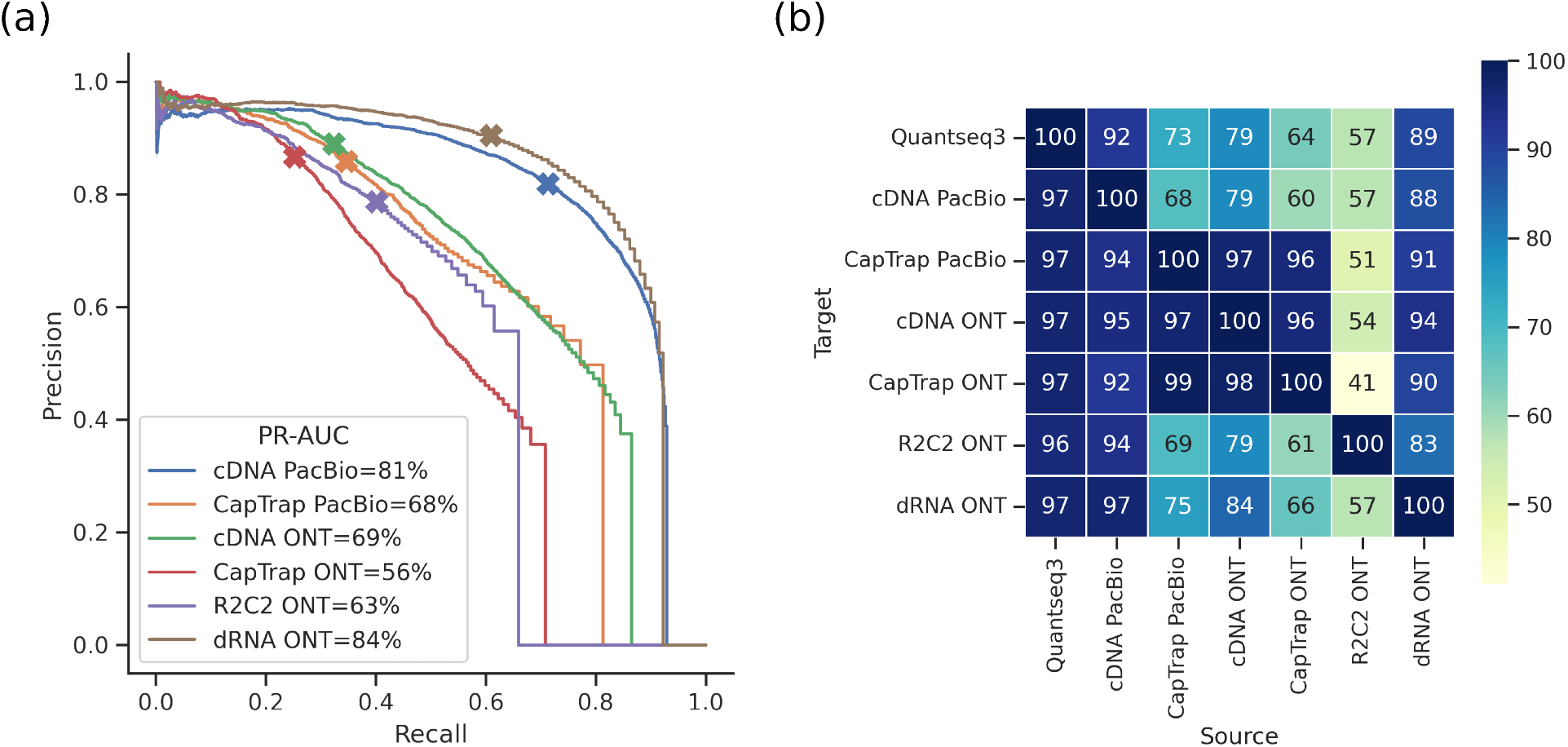
(a) Precision recall curve of LR-RNA-seq platforms/protocols against Quantseq3 ground truth. (b) Overlap of poly(A)-sites between different platforms/protocols.

### 3.4 Correction of transcript start and end sites

LAPA has a highly optimized strategy for transcript end site (TES) detection, and the same strategy can be adapted for the transcript start site (TSS). Therefore, we investigated if LAPA can improve the TESs and TSSs of the transcriptome annotation (GTF) generated with LR-RNA-seq tools. We ran TALON [35] on WTC11 samples and generated a GTF file per sample using WTC11 LR-RNA-seq samples. TALON compares the intron chain of detected transcripts against reference genome annotation and classifies transcript isoforms into categories of Known (full intron chain match with annotation), prefix/suffix ISM (incomplete splice match; introns are a subset of annotated intron chain), NIC (novel in catalog; intron chain contains novel intron with annotated donor and acceptor sites) NNC (novel not in catalog; intron chain contains a novel donor or acceptor site). TESs detected by TALON were compared against the Quantseq3 ground truth to obtain the percentage of supported TESs. We observed that only 55 to 70% of transcript end sites from TALON are supported by Quantseq3 across the platforms/protocols. Then, we corrected TESs from TALON using LAPA and re-calculated TES support (Figure-5.a-b, Supplementary Figure-6, Supplementary Table-8). TES correction with LAPA improves TES support by 25 ± 2% consistently across all samples. Similarly, we investigated if LAPA can be adapted for TSS correction. We compared TSSsagainst CAGE ground truth to obtain experimental support for TSSs (Figure-5.c-d, Supplementary Figure-7, Supplementary Table-9). Adapting the same correction strategy on TALON TSSss results in an improvement of 26 ± 9% across all the samples. The major improvement is obtained from the known category where TALON reports TSS or TES of transcripts with matching intron chains from reference genome annotation. Another improvement is observed in the ISM category, which is assumed to be partial transcript isoforms and usually discarded for downstream analysis. However, LAPA filters out partial transcripts but rescues some of the ISM transcripts that resulted from true TSS and TES in the internal exons. Internal terminal exons, also called hybrid-exons [9], have been underreported until recently, yet those exons are important to capture full tissue-specific transcriptome diversity. NIC and NNC transcripts already have a high level of CAGE and Quantseq3 support, but LAPA further increases the transcript diversity by reporting additional start and end sites for transcripts with the same intron in those categories. Overall, our benchmark results based on the transcript start and end sites indicate that LAPA can correct TSSs and TESs of transcriptome annotation generated with LR-RNA-seq tools; as a result, LAPA increases the transcript diversity results of alternative TSS and TES.

**Figure 5:**
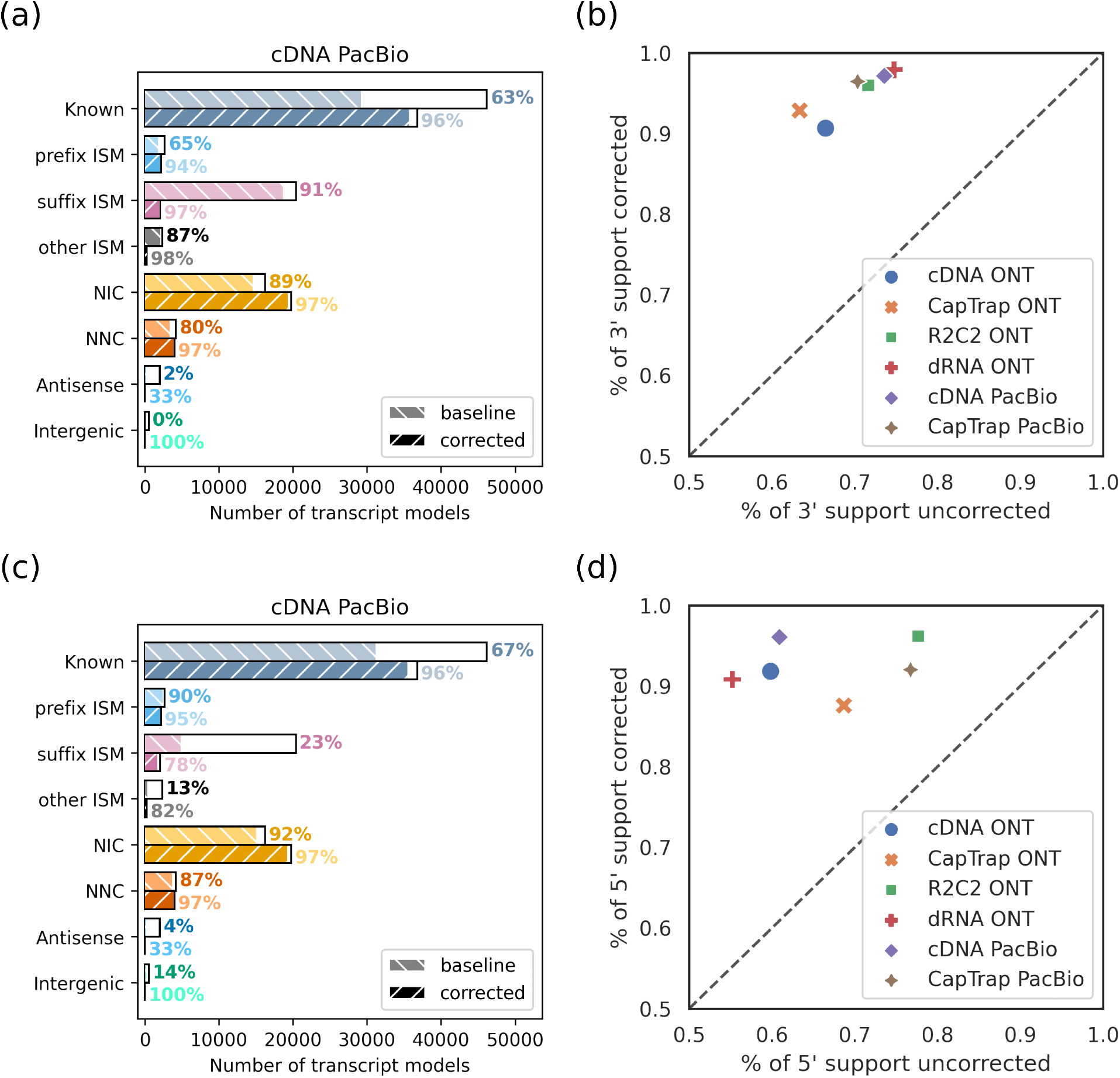
(a) Quantseq3 support of TES of transcript isoforms constructed from PacBio cDNA protocol with TALON before and after transcript end site correction with LAPA. (b) TES support of other protocols before and after correction. (c) CAGE support of TSS of transcript isoforms constructed from PacBio cDNA protocol. (d) TSS support of all platforms/protocols before and after correction.

### 3.5 Alternative polyadenylation during myogenesis

LAPA performs statistical testing to detect significant alternative polyadenylation events between conditions. As a case study, we investigated the signature of APA during C2C12 differentiation from myoblast cells to myotube cells [26]. We used two C2C12 undifferentiated myoblast replicates and two C2C12 3-day differentiated myotube replicates, both sequenced with PacBio LR-RNA-seq (Figure-6.a). We performed statistical testing between the myoblasts and myotubes with LAPA using Fisher’s exact test (Supplementary Methods-4,5). We detect unique signatures of poly(A)-usage between myoblast and myotube samples and observe high agreement between replicates (Figure-6.c). We detected 788 genes with significant alternative polyadenylation between the two conditions, and the top genes are annotated in the volcano plot figure-6.b (Supplementary Table-10). Many of these genes have previously reported biological significance in myogenesis. For example, *TPM1* is a myogenic factor and muscle structural protein [5]. *MATR3* is an RNA binding protein (RBP) that binds to myogenesis transcripts such as *MYOG* [2] and knockdown of *MATR3* impairs differentiation into mature myotubes. Tissue-specific APA signature of *MATR3* has also been previously reported [21]. Similarly, *LRRFIP1* knockdown reduces myoblast differentiation because *LRRFIP1* is a repressor of known muscle differentiation inhibitors [17, 33]. Similar experimental results are available, linking top significant reported genes to myogenesis: *NES (Nestin)* [37, 15], *Slc6a6 (Taut)* [32], *NPNT* [18], *PLEC* [1, 36]. *FSTL1* is an interesting example to highlight the relationship between miRNA binding and alternative polyadenylation. *Fstl1* is targeted by *miR-206*, which is a known MyomiR [29] and upregulated by *MyoD* [27]. We further investigated the relationship between gene expression and the 3’ UTR length of genes with significant alternative polyadenylation (Supplementary Figure-8). 3’ UTR shortening is significantly (*P* = 10^−68^) associated with higher gene expression, while 3’ UTR elongation is associated with lower gene expression (*P* = 10^−5^) compared to genes without significant APA (Figure-6.d). This difference in gene expression demonstrates the importance of regulatory elements such as miRNA binding sites in alternatively polyadenylated 3’ UTR region.

**Figure 6:**
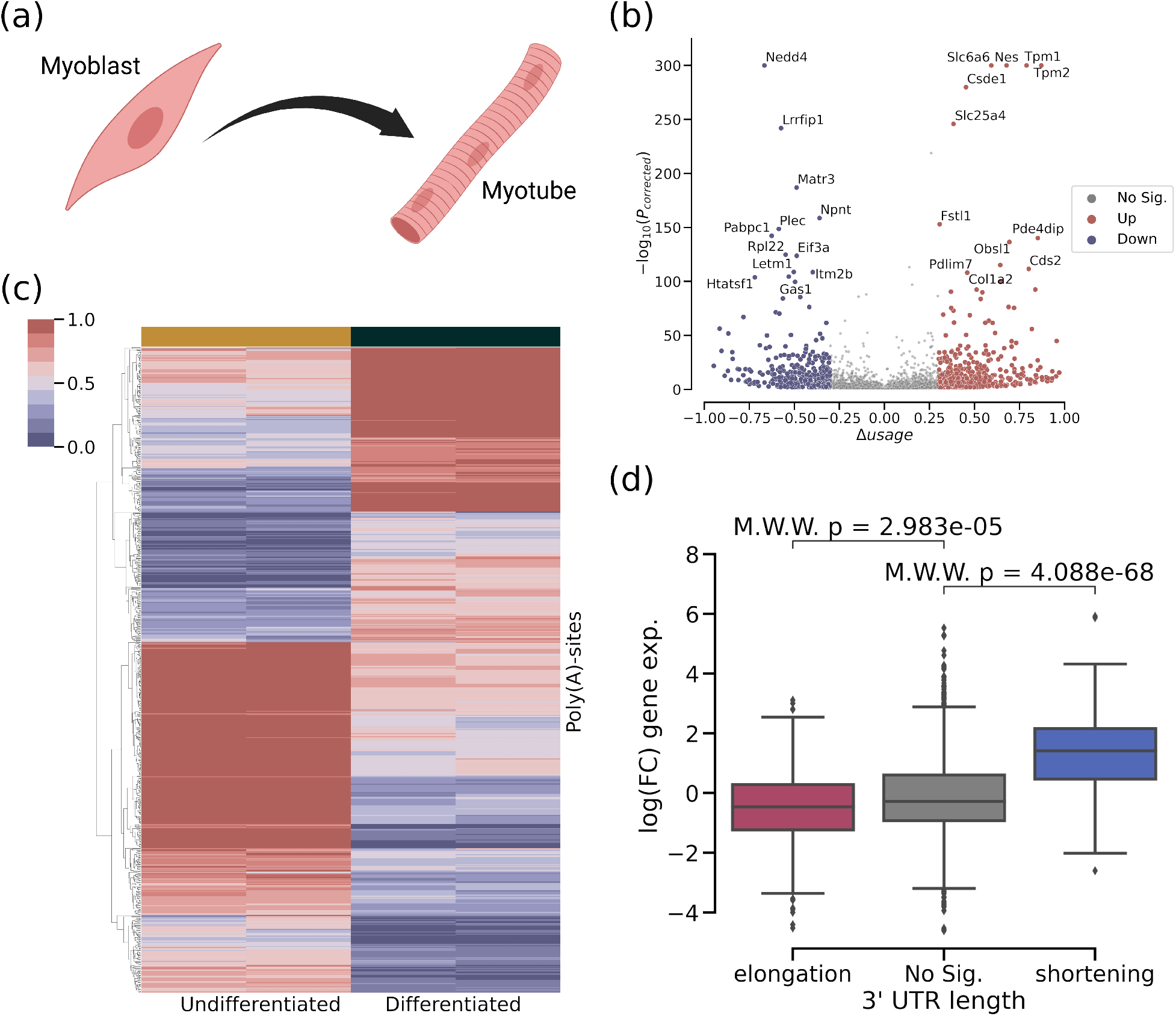
(a) Myogenesis (myoblast to myotube differentiation) (b) Volcano plot of genes with APA (c) Heatmap of poly(A)-usage metric for significant poly(A)-sites for undifferentiated myoblast and differentiated myotube samples (d) 3’ UTR length change during the differentiation against gene expression level.

## 4 Discussion

In this study, we introduced LAPA, a computational toolkit to analyze alternative polyadenylation using both long-read RNA-seq and 3’ sequencing. LAPA performs read-end counting, poly(A)-site clustering, peak-calling, annotation of poly(A)-sites based on the regulatory sequence elements, filters potential internal priming sites, and controls for replication rate. Our benchmark based on a comparison between independent platforms and protocols shows that LAPA detects APA with high precision and recall. Also, detected poly(A)-sites in expressed genes (>5 TPM) have a high level of poly(A)-signal support. Results presented in this paper demonstrate that LR-RNA-seq overcomes the limitation of short-read RNA-seq and enables accurate detection of polyadenylation sites independent of the platform/protocol. Yet, ONT dRNA and cDNA PacBio protocols have the best performance for poly(A)-site detection.

In addition to TES detection, we easily repurposed LAPA for TSS detection due to the modularity of the software. Using TSS and TES clusters detected by LAPA, transcriptome annotations generated from LR-RNA-seq tools can be further improved. During transcript correction, we rescue transcript isoforms previously annotated as partial transcripts (ISMs). A comparison of those rescued ISMs against the CAGE and Quantseq3 shows that those rescued transcripts have true TSS/TES in their internal exons. A similar comparison of TSS and TES of known transcript isoforms against the ground truth emphasized that the ends of transcripts in GENCODE annotation are not always accurate and can be improved with LR-RNA-seq using our proposed methodology. Overall, LAPA increases TSS/TES diversity of transcriptome, enabling downstream applications involving either end of transcripts, such as alternative promoter usage or miRNA binding.

LAPA performs statistical testing to detect APA. Our investigation of myoblast to myotube differentiation of C2C12 cell lines demonstrates the unique signature of APA during cell differentiation. Also, there is a significant association between APA and gene expression. Specifically, elongation of the 3’ UTR during the differentiation is associated with lower gene expression, while shortening is correlated with higher gene expression. Overall, our results demonstrate that alternative polyadenylation is essential for cell state/identity and regulation of gene expression.

## Availability and implementation

All the results in the paper are implemented in a snakemake [24] workflow format and reproducible with a single command.

**LAPA:** https://github.com/mortazavilab/lapa

**betabinomial:** https://github.com/MuhammedHasan/betabinomial

**gencode_utr_fix:** https://github.com/MuhammedHasan/gencode_utr_fix

## Author’s Contributions

M.H.C implemented the software and performed analysis, A.M. supervised the project, M.H.C and A.M. wrote the manuscript.

## Acknowledgements

We thank Fairlie Reese and Jasmine Sakr for their insightful comments on the manuscript.

