## Supplementary Material for "Analysis of alternative polyadenylation from long-read or short-read RNA-seq with LAPA"

*To whom correspondence should be addressed.

### Supplementary Figures

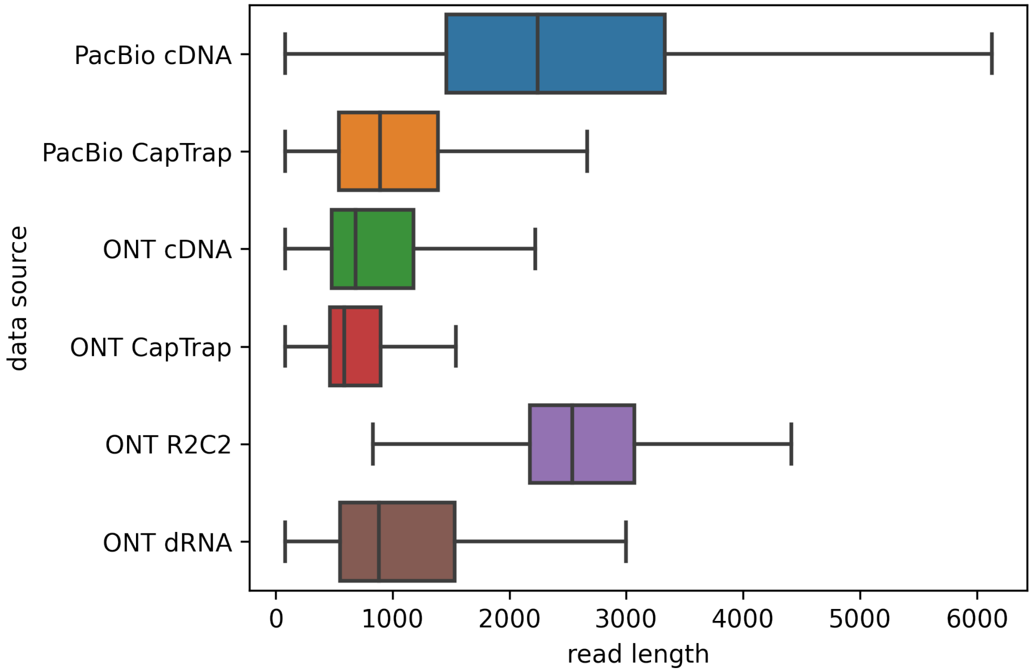


**Figure 1:** Read length distribution across platforms/protocols.

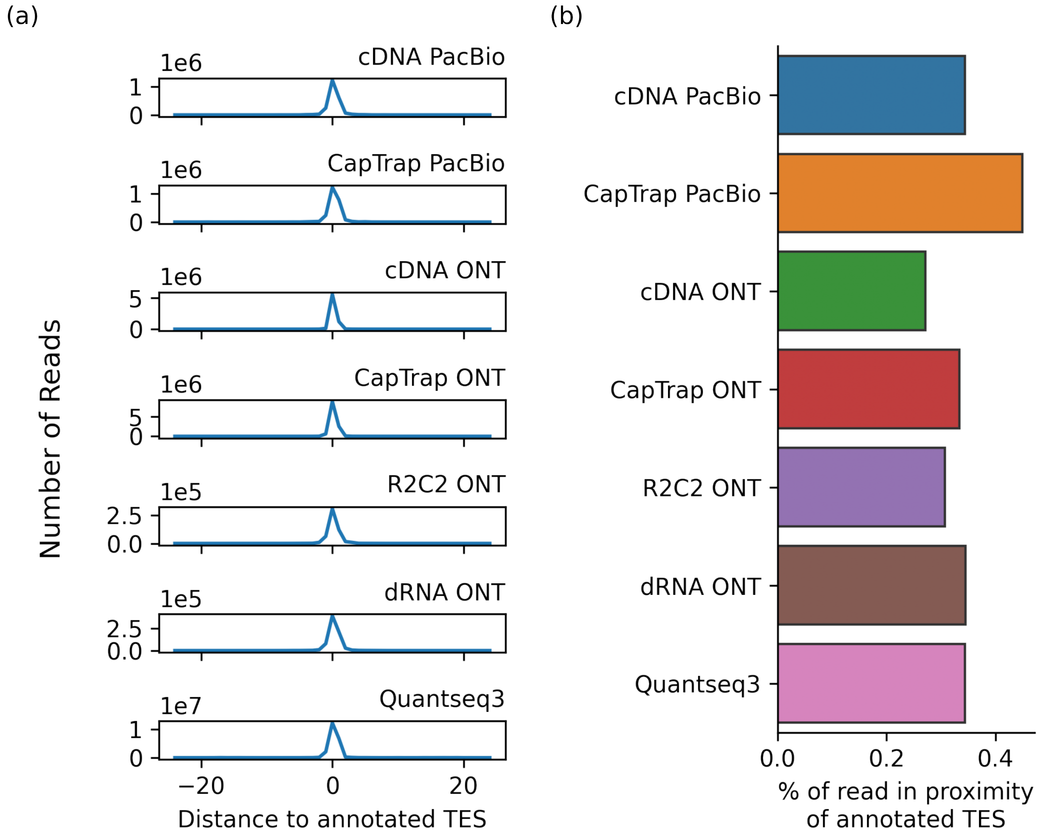


**Figure 2:** (a) Number of reads in the vicinity of annotated GENCODE transcript end site (TES). (b) Percentage of reads in the vicinity of annotated GENCODE TES.

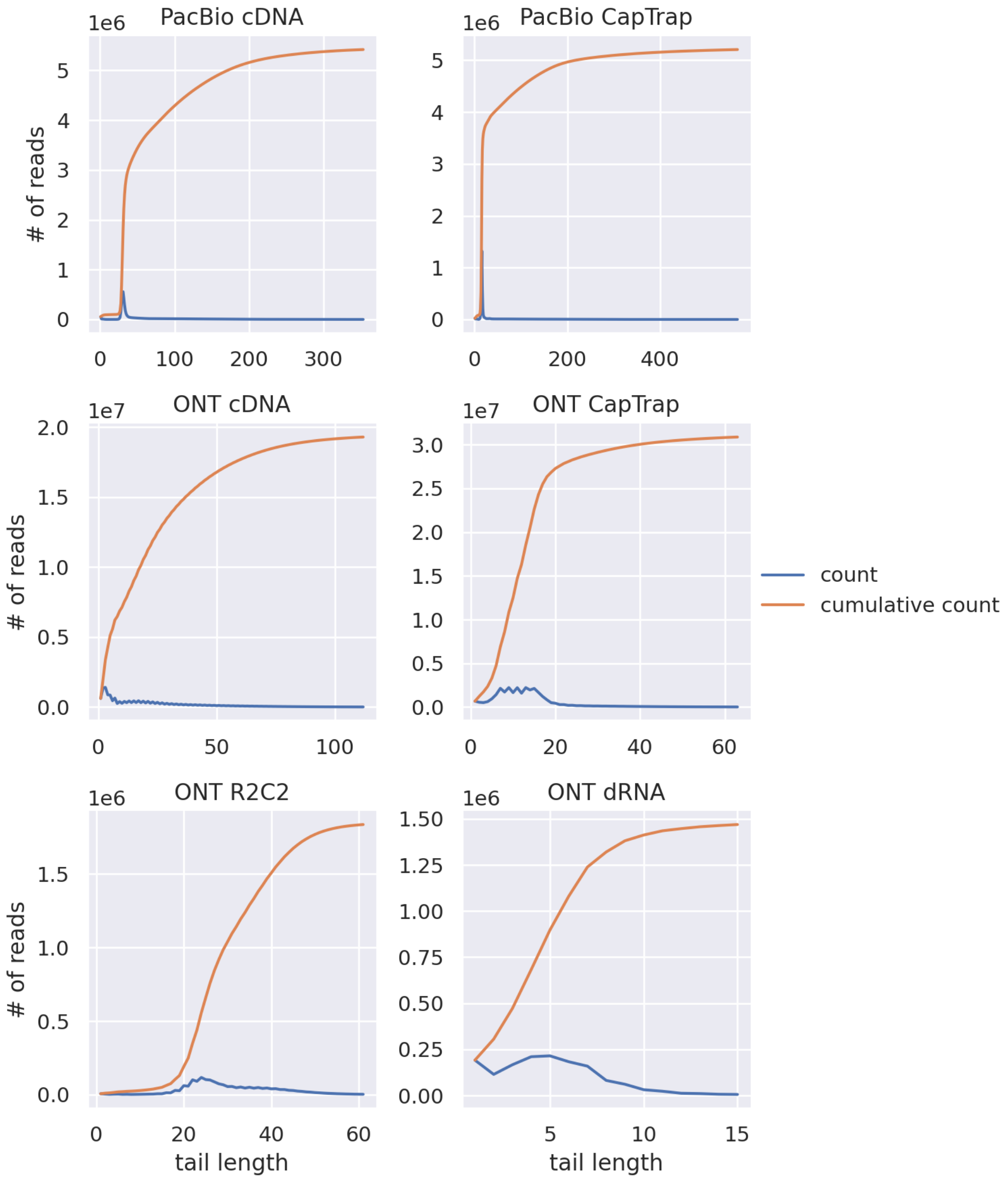


**Figure 3:** Poly(A)-tail length of reads across the platforms/protocols. Counts in blue and cumulative counts in orange. The distribution of counts indicates that poly(A)-tails of dRNA are trimmed, and poly(A)-tails of ONT cDNA and CapTrap reads are partially trimmed.

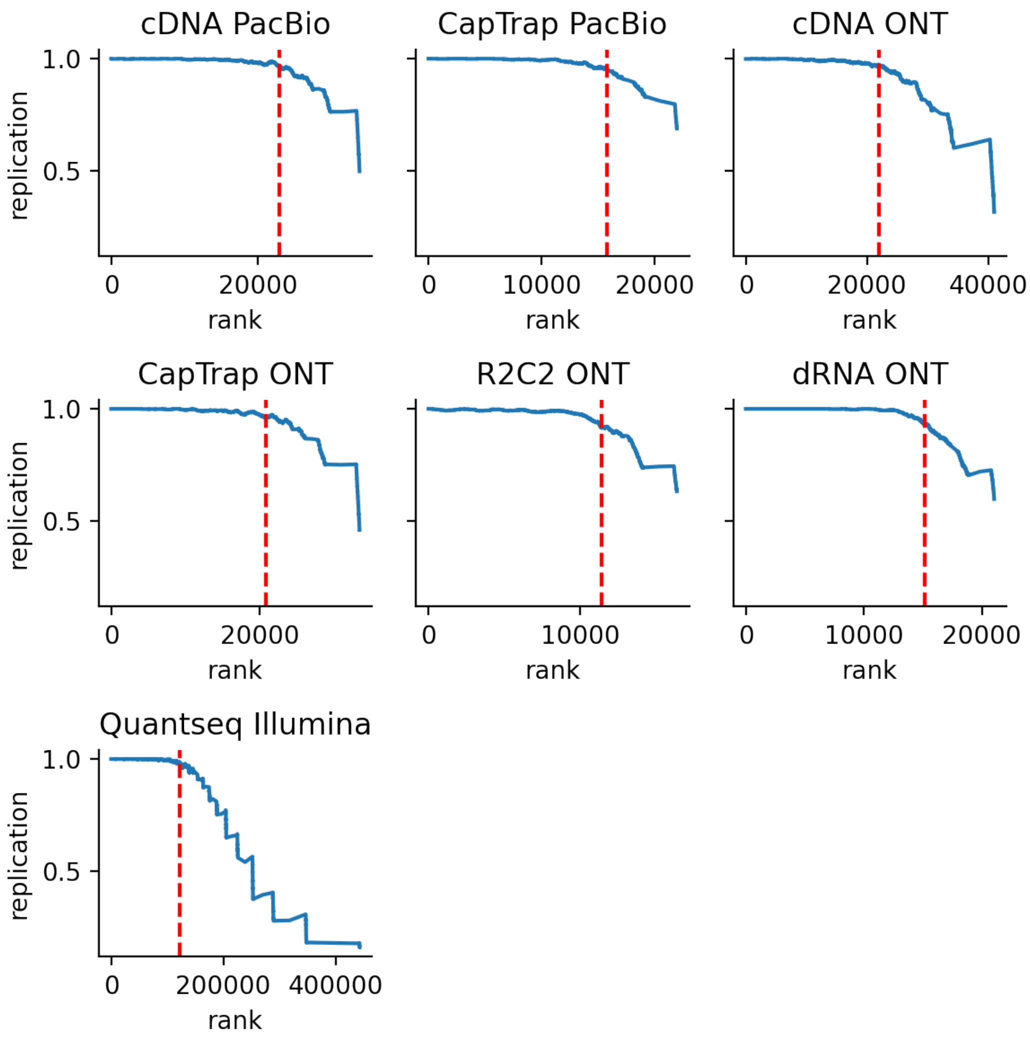


**Figure 4:** Replication rate of poly(A)-clusters between three experimental replicates. The red line represents a threshold for 95% replication rate. Quantseq3 produces the most replicated clusters at the same replication rate of 95% due to much deeper read coverage.

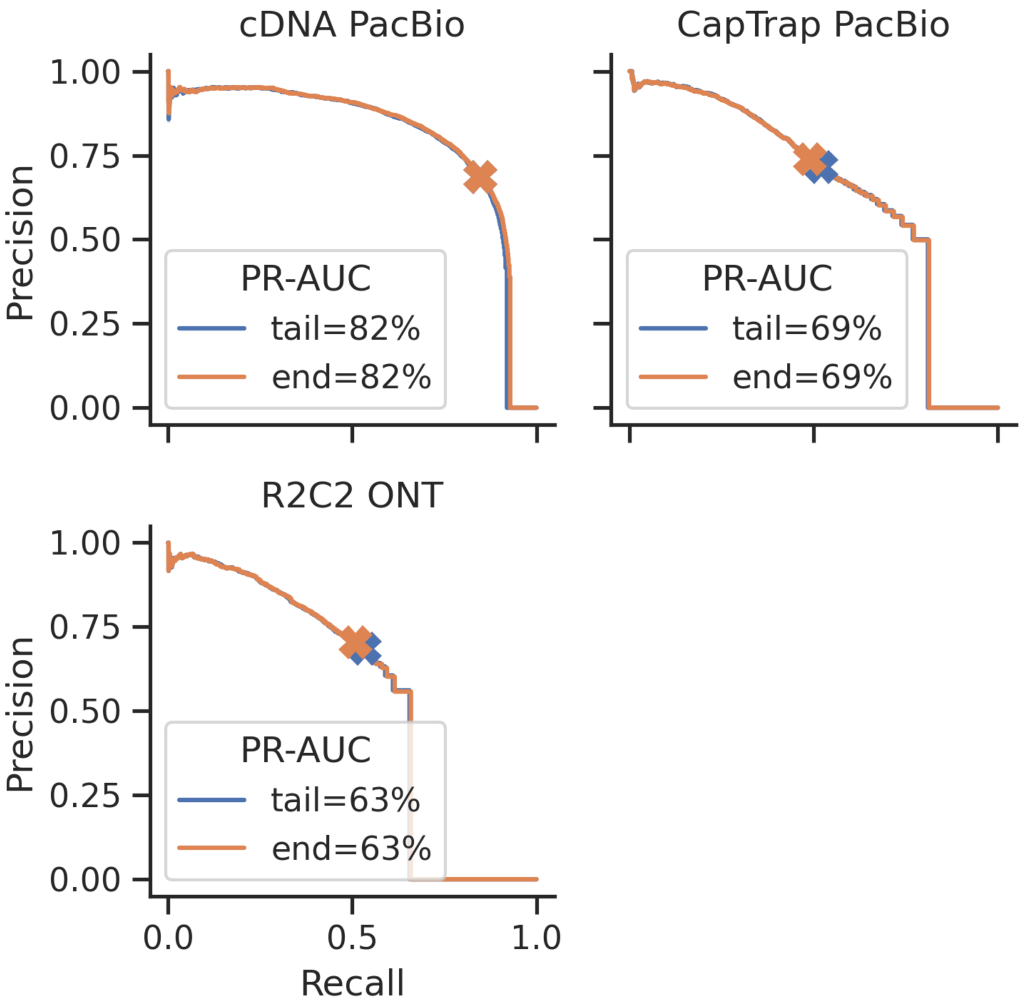


**Figure 5:** Effect of read-end counting parameters on precision-recall. Results show that requiring 20 bp poly(A)-tail has nearly no effect on the performance. Cross marks indicate the threshold based on 95% replication rate.

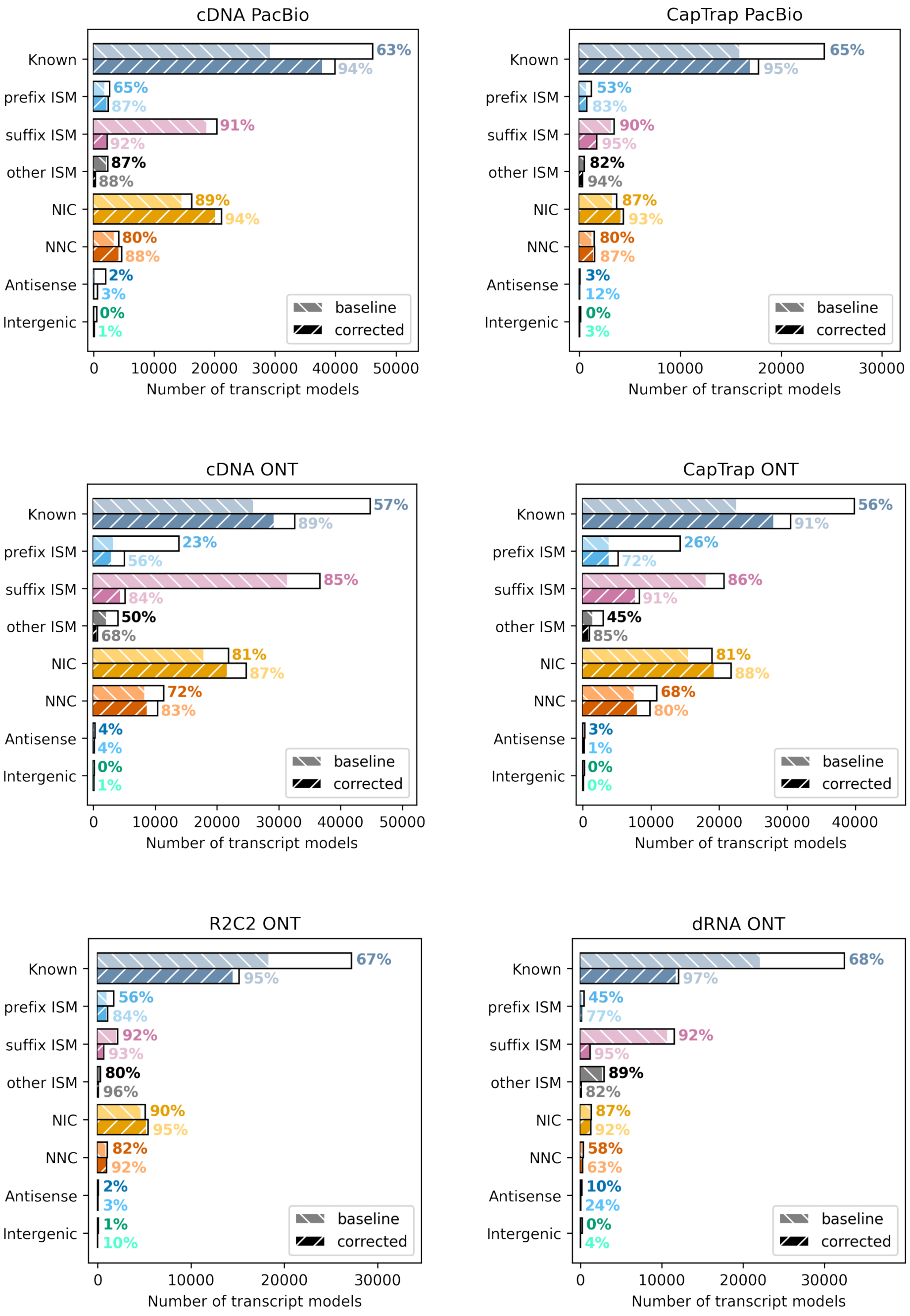


**Figure 6:** Quantseq3 support of TES of transcript isoforms constructed with TALON before and after transcript end site correction with LAPA.

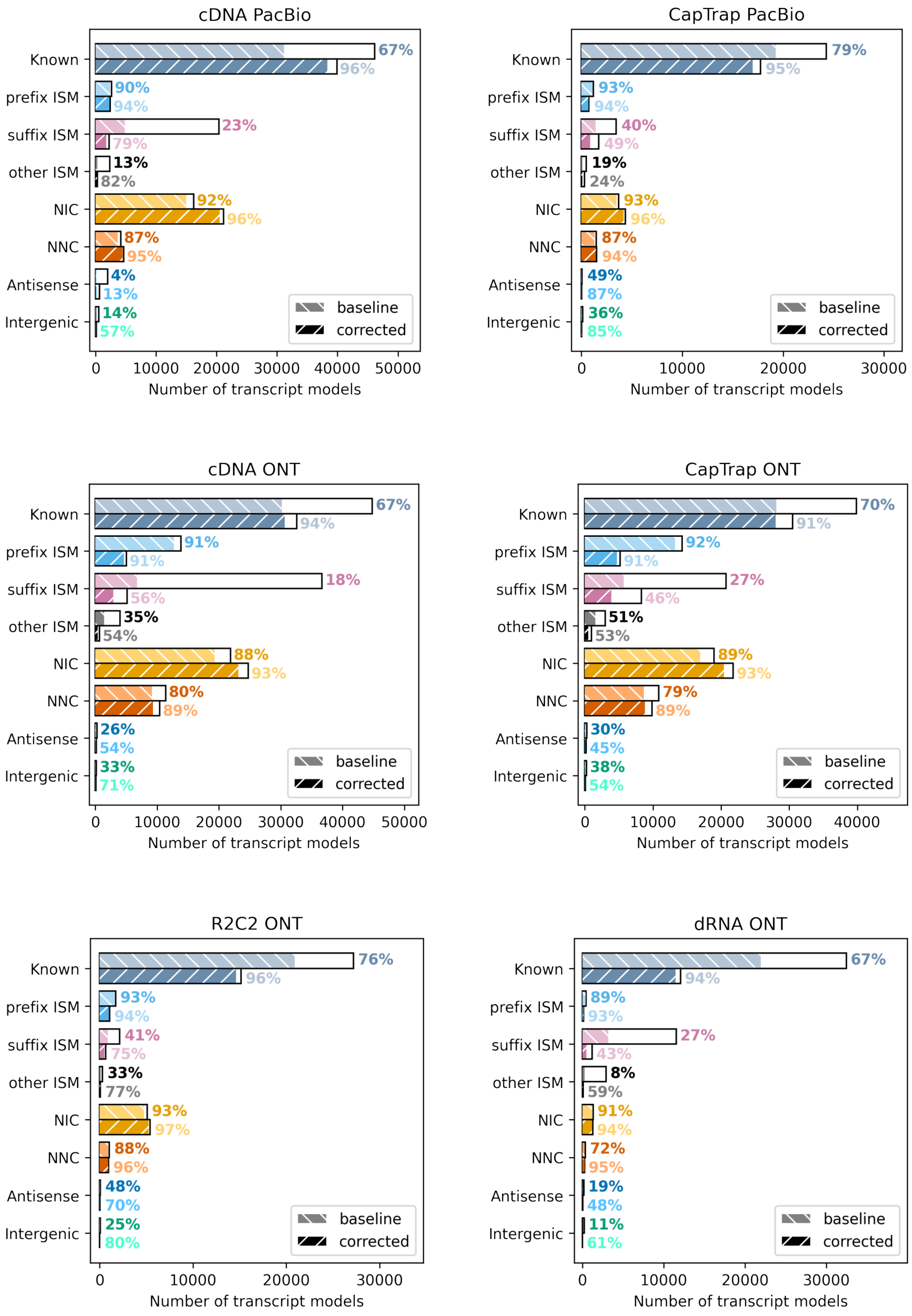


**Figure 7:** CAGE support of TSS of transcript isoforms constructed with TALON before and after transcript start site correction with LAPA.

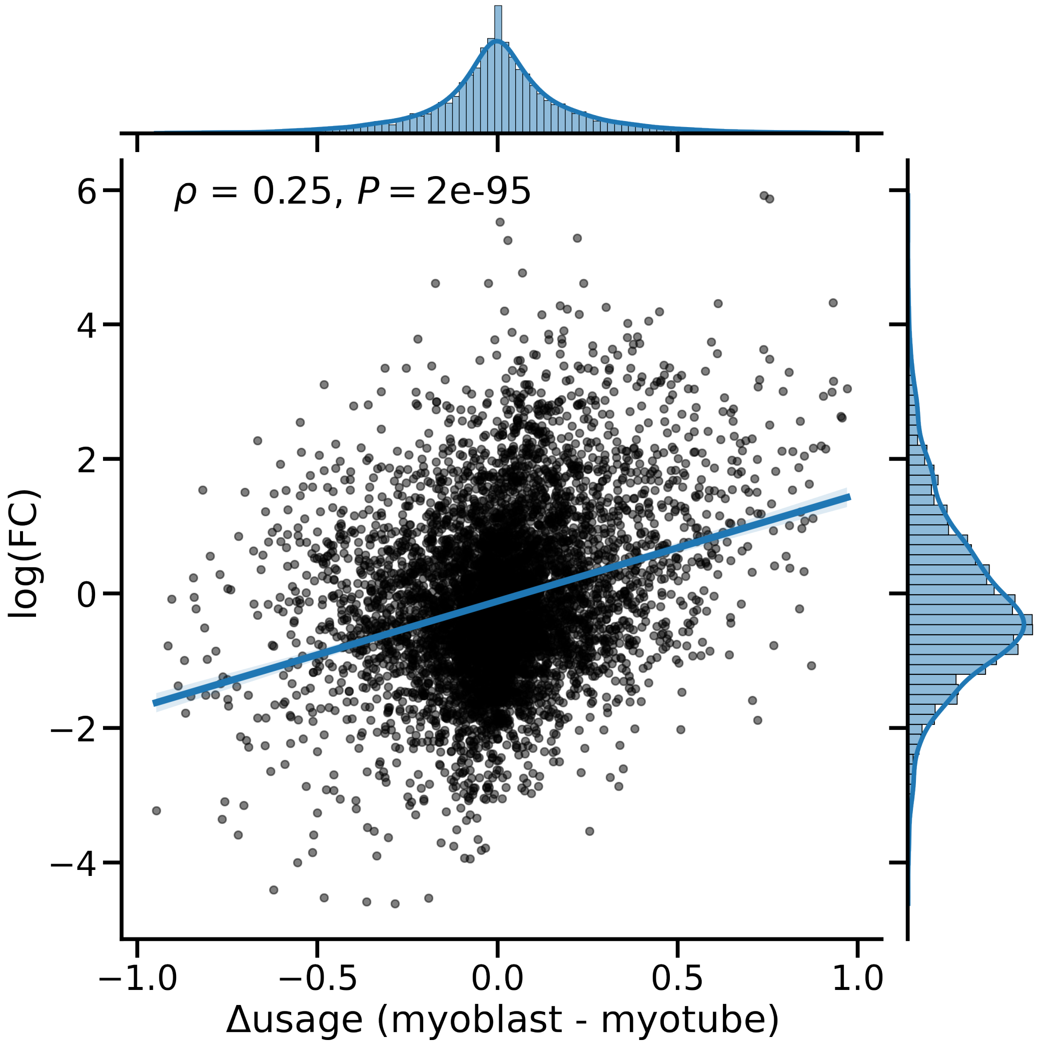


**Figure 8:** Changes in distal poly(A)-site usage during myoblast to myotubes differentiation correlates with gene expression log-fold change between conditions.

### Supplementary Tables

**Supplementary Table-1:**

| data source | read | count |
|---|---|---|
| Illumina RNA-seq | All Reads | 34,940,279 ± 3,565,959 |
| Illumina RNA-seq | Poly(A) usable reads | 66,276 ± 9,391 |
| ONT CapTrap | All Reads | 16,967,434 ± 2,732,784 |
| ONT CapTrap | Poly(A) usable reads | 12,706,021 ± 2,141,450 |
| ONT R2C2 | All Reads | 663,342 ± 110,955 |
| ONT R2C2 | Poly(A) usable reads | 656,352 ± 110,504 |
| ONT cDNA | All Reads | 17,064,845 ± 6,345,987 |
| ONT cDNA | Poly(A) usable reads | 9,144,553 ± 3,696,101 |
| ONT dRNA | All Reads | 996,143 ± 722,993 |
| ONT dRNA | Poly(A) usable reads | 902,966 ± 653,236 |
| PacBio CapTrap | All Reads | 2,133,211 ± 340,192 |
| PacBio CapTrap | Poly(A) usable reads | 1,906,174 ± 323,949 |
| PacBio cDNA | All Reads | 2,474,974 ± 698,776 |
| PacBio cDNA | Poly(A) usable reads | 2,373,264 ± 703,259 |
| Quantseq3 | All Reads | 105,423,691 ± 16,713,839 |
| Quantseq3 | Poly(A) usable reads | 26,515,255 ± 3,443,010 |

**Supplementary Table-2:**

| data source | count | mean | std | min | 25% | 50% | 75% | max |
|---|---|---|---|---|---|---|---|---|
| ONT CapTrap | 38118062 | 738 | 418 | 80 | 465 | 587 | 897 | 8327 |
| ONT R2C2 | 1984417 | 2701 | 769 | 80 | 2174 | 2536 | 3070 | 9001 |
| ONT cDNA | 27433659 | 912 | 647 | 80 | 482 | 682 | 1179 | 11007 |
| ONT dRNA | 2730704 | 1200 | 966 | 81 | 551 | 884 | 1530 | 16744 |
| PacBio CapTrap | 5773371 | 1042 | 596 | 80 | 539 | 894 | 1390 | 7518 |
| PacBio cDNA | 7147240 | 2525 | 1482 | 80 | 1463 | 2241 | 3329 | 20313 |

**Supplementary Table-3:**

| Data Source | median_tail_length |
|---|---|
| PacBio cDNA | 35 |
| PacBio CapTrap | 16 |
| ONT cDNA | 17 |
| ONT CapTrap | 12 |
| ONT R2C2 | 29 |
| ONT dRNA | 5 |

**Supplementary Table-4:**

| three_prime_utr | intron | exon | Data Source |
|---|---|---|---|
| 0.976678 | 0.017077 | 0.006245 | cDNA PacBio |
| 0.955889 | 0.025308 | 0.018804 | CapTrap nPacBio |
| 0.949037 | 0.028951 | 0.022012 | cDNA ONT |
| 0.891419 | 0.065609 | 0.042971 | CapTrap ONT |
| 0.982239 | 0.009055 | 0.008706 | R2C2 ONT |
| 0.987273 | 0.007879 | 0.004848 | dRNA ONT |
| 0.876626 | 0.096488 | 0.026885 | Quantseq Illumina |

**Supplementary Table-5:**

| data source | % of read in proximity of annotated TES |
|---|---|
| cDNA PacBio | 0.346173 |
| CapTrap PacBio | 0.457267 |
| cDNA ONT | 0.273562 |
| CapTrap ONT | 0.336798 |
| R2C2 ONT | 0.305878 |
| dRNA ONT | 0.342175 |
| Quantseq3 | 0.345066 |

**Supplementary Table-6:**

| Data Source | threshold |
|---|---|
| cDNA PacBio | 6 |
| CapTrap PacBio | 5 |
| cDNA ONT | 9 |
| CapTrap ONT | 5 |
| R2C2 ONT | 6 |
| dRNA ONT | 6 |
| Quantseq Illumina | 12 |

**Supplementary Table-7:**

| Recall | Precision | Data Source |
|---|---|---|
| 0.852165 | 0.682092 | cDNA PacBio |
| 0.489438 | 0.738593 | CapTrap PacBio |
| 0.406448 | 0.833830 | cDNA ONT |
| 0.355482 | 0.746878 | CapTrap ONT |
| 0.513217 | 0.698884 | R2C2 ONT |
| 0.813993 | 0.786981 | dRNA ONT |

**Supplementary Table-8:**

| Uncorrected | Corrected | Data Source |
|---|---|---|
| 0.598391 | 0.918254 | cDNA ONT |
| 0.686895 | 0.875743 | CapTrap ONT |
| 0.776502 | 0.962058 | R2C2 ONT |
| 0.552042 | 0.908499 | dRNA ONT |
| 0.609074 | 0.960579 | cDNA PacBio |
| 0.767461 | 0.920187 | CapTrap PacBio |

**Supplementary Table-9:**

| Uncorrected | Corrected | Data Source |
|---|---|---|
| 0.664829 | 0.906510 | cDNA ONT |
| 0.633491 | 0.928476 | CapTrap ONT |
| 0.716032 | 0.959905 | R2C2 ONT |
| 0.747163 | 0.979217 | dRNA ONT |
| 0.735696 | 0.971293 | cDNA PacBio |
| 0.703576 | 0.964382 | CapTrap PacBio |

### Supplementary Material

#### 1 - Platform/Protocols-specific preprocessing of raw data

In this project, we utilized the publicly available LRGASP dataset available on ENCODE portal. Fastq files are used to count a raw number of reads generated by the protocols/platforms, and bam files are used for the remaining downstream analysis. Reference genome sequence and annotation downloaded from GENCODE and genome version of GRCh38 (v.38) for human and GRCm38 (vM25) for mouse are used.

**Quantseq3:** We downloaded fastq and bam files for Quantseq3 with ENCODE identification numbers ENCFF034QBE, ENCFF544FMU, ENCFF201HMB and ENCFF399HRH, ENCFF364WSF, ENCFF146APZ respectively. Bam files were previously processed by seqkit [7] to subset reads with 18 bp poly(A)-tails, bbduk.sh [2] to trim adapter sequences and finally aligned to genome with STAR [3] with recommended arguments.

**Short-read RNA-seq:** We downloaded fastq files for single-ended short read RNA-seq with ENCODE identification numbers ENCFF492PIB, ENCFF365HXR, ENCFF621NOL, ENCFF738DHG, ENCFF235BRN, ENCFF742FDJ. Then we subset the reads with at least 18 bp poly(A)-tails and aligned them to the genome with a STAR aligner.

Long-read RNA-seq protocols are aligned with minimap2 [4] with recommended arguments for each protocol and pipeline. Gene annotation is provided to minimap2 using paftools.js tools to prioritize known splice sites.

**PacBio:** We downloaded fastq files for PacBio cDNA (ENCFF563QZR, ENCFF370NFS, ENCFF245IPA) and CapTrap (ENCFF105WIJ, ENCFF212HLP, ENCFF003QZT). Then, fastq files are aligned to the reference genome using minimap2 with the high-quality flag ('-ax splice:hq').

**ONT:** We downloaded fastq files for ONT cDNA (ENCFF263YFG, ENCFF023EXJ, ENCFF961HLO) and CapTrap (ENCFF654SNK, ENCFF934KDM, ENCFF104BNW). Those two protocols, unlike other protocols, are not stranded; thus, we disabled the forward strand argument ('-uf'). In this setting, minimap2 aligns reads to the best matching strand and annotates strand information with 'ts' flag in the alignment file based on splicing dinucleotides (GT-AG); yet preserves the original direction of sequence in the alignment. Thus, we wrote to custom script to reserve reads in the complementary strand based on 'ts' flag to obtain reads in the correct strand. Fastq files for

ONT dRNA (ENCFF155CFF, ENCFF771DIX, ENCFF600LIU) and R2C2 (ENCFF153SIE, ENCFF377IEH, ENCFF489PQQ) are downloaded. In dRNA alignment '-k14' argument, which is recommended for noise dRNA reads, is used. R2C2 is aligned with the default argument. In ONT protocols, we applied just '-ax splice' argument without a high-quality parameter ('hq') because ONT reads are noisier than PacBio reads.

#### 2 - LR-RNA-seq base-calling and poly(A)-tail trimming

In this study, we did not perform base-calling on LR-RNA-seq samples but rather downloaded base-called fastq files. However, base-calling pipelines (Isoseq3 for PacBio and guppy for ONT) trim poly(A)-tails by recommended default; thus, base-calling affects the polyadenlation related downstream analysis. In PacBio base-calling, Isoseq3 has an argument called '–require-polya', which detects the specified number of A base pairs at the 3' end of the read and filters reads lacking a poly(A)-tail. However, this argument also trims the poly(A)-tail sequence resulting in final reads with trimmed poly(A)-tails. PacBio datasets we used in this study are not filtered with '–require-polya' argument, so PacBio reads have poly(A)-tail sequence at the 3' end of the reads (Supplementary Figure-3). GUPPY is software to perform base-calling on ONT platform, which trims adaptors and poly(A)-tail sequence by default on RNA basecalling. GUPPY users need to set '–trim_strategy None' to preserve poly(A)-tails [6]. Poly(A)-tails are trimmed in ONT dRNA samples used in the study. In ONT cDNA and CapTrap protocols, poly(A)-tails in the forward strand are trimmed while poly(A)-tail of reads in the complementary strand are not trimmed. Because those samples are not stranded, GUPPY is only able to detect A base pairs at the 3' end of the and kept T bases in the 5' end of the reads in the complementary strand. Poly(A)-tails in ONT R2C2 reads are not trimmed, and most of the reads have at least 20 bp of poly(A)-tail (Supplementary Figure-3).

#### 3 - The output file structure of LAPA and content of files

```
example_output_dir
├── polyA_clusters.bed
├── raw_polyA_clusters.bed
├── counts
│   ├── all_polyA_counts_neg.bw
│   ├── all_polyA_counts_pos.bw
│   ├── {sample}_polyA_counts_neg.bw
│   └── {sample}_polyA_counts_pos.bw
├── coverage
│   ├── all_polyA_coverage_neg.bw
│   ├── all_polyA_coverage_pos.bw
│   ├── {sample}_polyA_coverage_neg.bw
│   └── {sample}_polyA_coverage_pos.bw
├── ratio
│   ├── all_polyA_ratio_neg.bw
│   ├── all_polyA_ratio_pos.bw
│   ├── {sample}_polyA_ratio_neg.bw
│   └── {sample}_polyA_ratio_pos.bw
├── dataset
│   └── {dataset}.bed
├── raw_sample
│   └── {sample}.bed
└── logs
    ├── final_stats.log
    ├── progress.log
    └── warnings.log
```

**polyA_clusters.bed:** is the main output of LAPA and contains poly(A) clusters with replication rate. Those set of clusters is a high confidence set of poly(A) clusters. Poly(A)-cluster bed file contains the following columns:

- **Chromosome:** Chromosome of the poly(A) cluster.
- **Start:** Start position of poly(A) cluster.
- **End:** End position of poly(A) cluster.
- **polyA_site:** Exact poly(A) site (peak) of the cluster (0-based).

- **count:** number of reads supporting to cluster (ending in the cluster).
- **Strand:** Strand of the poly(A) cluster.
- **Feature:** Genomics feature overlapping with the cluster (obtained from GTF file).
- **gene_id:** The gene containing the poly(A) clusters.
- **tpm:** TPM of the cluster calculated by $\frac{count*1{,}000{,}000}{\sum count}$.
- **gene_count:** Total number reads in all the clusters of this gene calculated by $\sum count_i$ where $i \in gene$.
- **usage:** Percentage use of specific poly(A) clusters of the gene calculated by $\frac{count}{gene\_count}$
- **fracA:** Number of A bp in the reference genome sequence in following 10 bp after the poly(A)-site.
- **signal:** Poly(A)-signal sequence detected in the vicinity of poly(A)-site. Annotated as 'position@signal' where the position is the position in the reference genome and signal indicates 6 bp poly(A)-signal sequence.
- **annotated_site:** End position of annotated poly(A)-site in 3' UTR based on the GTF if poly(A) cluster located in 3' UTR.

**raw_polyA_clusters.bed:** contains all the poly(A) clusters detected by LAPA so not filtered for replication. counts: is a directory containing read end bigwig files. Each bigwig file contains a number of read-end counts per position indicating possible poly(A)-sites. This directory contains one bigwig file for each strand and is not filtered, so it represents raw data. There are pairs of bigwig files per sample per strand. The file starting with "all" prefixes contains counts from all the samples where total counts are aggregated into one bigwig file by summation. coverage: is a directory containing bigwig files for coverage. Each file contains coverage of each position with non-zero values in the read-end count file. So the file format is sparse, has values only for positions where at least 1 read is ending, and the remaining positions are zero despite coverage can be non-zero. A sparse file format is used to limit file size and increase computational efficiency. The file starting all prefixes contains counts from all the samples where counts are aggregated into one bigwig file. ratio: is a directory containing a ratio of read-end counts to coverage (). This ratio indicates percentage reads are ending at a position given coverage of the position. If the ratio is close to one and there is high coverage, the site is definitive poly(A)-site. If the ratio is close to 0, then the reads could be ending at the position by chance. Based on the default parameters LAPA (cluster_ratio_cutoff) only initialize cluster if ratio > 5% at a position. The file starting all prefixes contains the ratio for all the samples.

**dataset:** is a directory containing poly(A)-cluster .bed files per dataset. Those files are filtered for replication rate using samples of the dataset, then replicated clusters from all the samples aggregate into the .bed file for the dataset.

**raw_sample:** is a directory containing poly(A) cluster .bed files per sample where files are not filtered for replication.

**sample:** is a directory containing poly(A) cluster .bed files per sample where files are filtered for replication.

**logs:** is a directory containing logs of LAPA. “final_stats.log” contains statistics about poly(A) clusters after the program is finished. “progress.log” provide inside about the progress of the program run. “warnings.log” file contains any possible warning encounters during the run time.

#### 4 - Statistical testing

**Fisher's exact test:** We perform fisher's exact test by contracting 2 x 2 contingency tables below:

| | |
|---|---|
| $c_0^i$ | $c_1^i$ |
| $\sum_j c_0^j - c_0^i$ | $\sum_j c_1^j - c_1^i$ |

where $c_0^i$ is the read-end count for a poly(A)-site in condition-0 and $c_1^i$ is read-end count for condition-1. $\sum_j c_0^j - c_0^i$ and $\sum_j c_1^j - c_1^i$ is the total read-end count of all poly(A)-sites in the gene subtracted from read-end count of poly(A)-site in question. Then, adjusted p-values of all poly(A)-sites are calculated using Benjamini/Hochberg multiple testing corrections method [1]. In addition to p-value and adjusted p-value, we report $\Delta Usage$ and odds-ratio between conditions for each poly(A)-site calculated by:

$$\frac{c_0^i / c_1^i}{(\sum_j c_0^j - c_0^i)/(\sum_j c_1^j - c_1^i)}$$

**Beta-binomial test:** To perform beta-binomail test, we first infer $\alpha_i$ and $\beta_i$ dispersion parameters for each poly(A)-site $i$ using the method of moments estimation [8]. There used to be no python package to perform fast vectorial method of moments estimation for beta-binomial; thus, we developed python package (github.com/muhammedhasan/betabinomial) to perform fast vectorial parameter inference and statistical testing with beta-binomial test. P-values based on test calculated by:

$$P_i^j = 2 * min(0.5, CDF(k_i^j | n_i^j, \alpha_j, \beta_j), 1 - CDF(k | n, \alpha_j, \beta_j))$$

where $k_i^j$ is read-end count of poly(A)-site $j$ in sample $i$, $n_i^j$ is total read-end count of gene poly(A)-site belong, CDF is the cumulative distribution function of BetaBinomial and $\alpha_j$, $\beta_j$ is dispersion parameters of the poly(A)-site. We calculate $\Delta Usage$ for each poly(A)-site with $Usage_{mean}^j - Usage_i^j$ where $Usage_{mean}^j$ calculated by $\frac{\alpha_j}{\alpha_j + \beta_j}$. Also, observed and expected read-end count ($\frac{n * \alpha_j}{\alpha_j + \beta_j}$) are reported. We further report log fold change based on the expected read-end counts:

$$logFC = \log \frac{k_i^j}{n * \alpha_j/(\alpha_j + \beta_j)}$$

and z-score statistics for each poly(A)-site:

$$Z_{score} = \frac{k_i^j - n * \alpha_j/(\alpha_j + \beta_j)}{\sqrt{n\pi(1-\pi)(1+(n-1)\rho)}}$$

$$\rho = \frac{1}{\alpha_i + \beta_i + 1}$$

$$\pi = \frac{\alpha_i}{\alpha_i + \beta_i}$$

#### 5 - Alternative polyadenylation during myogenesis

Two experimental replicates of myoblast bam files with identifiers of ENCFF772LYG and ENCFF421MIL myotube files with ENCFF699KOR and ENCFF731HHB are downloaded from ENCODE portal. We run LAPA with its default arguments on the samples and replicates used to filter poly(A)-cluster to ensure 95% replication rate. Based on fisher's exact test, p-values are obtained and correct for familywise false discovery rate using Benjamini/Hochberg multiple testing corrections method [1]. Significant poly(A)-sites has $P < 0$ and $|\Delta usage > 0.3|$. For downstream analysis in figure-6.b-c, we subset the most significant poly(A)-site in each gene. We annotated upregulation and downregulation of poly(A)-site based on the sign of $\Delta usage > 0.3$; however, the sign of $\Delta usage$ does not matter in gene-level given alternative polyadenylation is a zero-sum game (poly(A)-usage of all poly(A)-sites in a gene sum up to 1). In figure-6.d, we subset the distal poly(A)-sites in the significant genes and annotated elongation and shorting of 3' UTR based on the sign of $\Delta usage$ distal poly(A)-site. Then, genes are categorized into elongation, shorting, and No sig. (no significance) and compared gene expression log-fold change between categories based on Mann Whitney U test [5]. We also plotted raw gene expression fold change between $\Delta usage$ distal poly(A)-site as a scatterplot in supplementary figure-8 without categorization. There is a significant correlation between 3' UTR shortening and gene expression with a Spearman's correlation coefficient of 0.25.
